# Predicting trait phenotypes from knowledge of the topology of gene networks

**DOI:** 10.1101/2021.06.29.450449

**Authors:** Andy Beatty, Christopher R. Winkler, Thomas Hagen, Mark Cooper

## Abstract

In many fields there is interest in manipulating genes and gene networks to realize improved trait phenotypes. The practicality of doing so, however, requires accepted theory on the properties of gene networks that is well-tested by empirical results. The extension of quantitative genetics to include models that incorporate properties of gene networks expands the long tradition of studying epistasis resulting from gene-gene interactions. Here we consider *NK* models of gene networks by applying concepts from graph theory and Boolean logic theory, motivated by a desire to model the parameters that influence predictive skill for trait phenotypes under the control of gene networks; *N* defines the number of graph nodes, the number of genes in the network, and *K* defines the number of edges per node in the graph, representing the gene-gene interactions. We define and consider the attractor period of an *NK* network as an emergent trait phenotype for our purposes. A long-standing theoretical treatment of the dynamical properties of random Boolean networks suggests a transition from long to short attractor periods as a function of the average node degree *K* and the bias probability *P* in the applied Boolean rules. In this paper we investigate the appropriateness of this theory for predicting trait phenotypes on random and real microorganism networks through numerical simulation. We show that: (i) the transition zone between long and short attractor periods depends on the number of network nodes for random networks; (ii) networks derived from metabolic reaction data on microorganisms also show a transition from long to short attractor periods, but at higher values of the bias probability than in random networks with similar numbers of network nodes and average node degree; (iii) the distribution of phenotypes measured on microorganism networks shows more variation than random networks when the bias probability in the Boolean rules is above 0.75; and (iv) the topological structure of networks built from metabolic reaction data is not random, being best approximated, in a statistical sense, by a lognormal distribution. The implications of these results for predicting trait phenotypes where the genetic architecture of a trait is a gene network are discussed.

## INTRODUCTION

The linear statistical model framework of quantitative genetics (Fisher 1918; Cockerham 1954; Kempthorne 1954; Henderson 1975; Falconer and Mackay 1996; Hill *et al.* 2008; Lynch and Walsh 1998; Barton *et al.* 2017; Walsh and Lynch 2018) has provided a means with which to explain heritable trait variation through the partitioning of genetic variance into additive and non-additive components and has been practically applied with great success by plant and animal breeders (Hallauer and Miranda 1988; Hill and Mackay 1989; Bernardo 2002; Meuwissen et al. 2001; Cooper *et al.* 2014; Rogers *et al.* 2021). Plant breeding procedures grounded within this quantitative genetics framework, for example, have contributed to a tripling of grain yield of maize in the United States since 1960 (Bruce *et al.* 2002; Duvick 2001).

While quantitative genetics theory has been used to great effect without the need for a detailed understanding of how specific genes work to determine trait phenotypes, the past few decades have brought a wealth of knowledge about the molecular structure and function of genes and proteins influencing quantitative traits, e.g., in the crop plant maize (*Zea mays*) (Simmons et al. 2021). Increasingly this research has shown that molecular components at work in the living cell interact with each other, often in highly specific and context-dependent networks (Holland 2001; Holland 2006; Benfey and MITCHELL-OLDS 2008; Marjoram *et al.* 2014). Davidson *et al.* (2002), for example, identified a genomic regulatory network of fifty genes controlling the temporal and spatial development of the endomesoderm in the sea urchin (*Strongylocentrotus purpuratus*) embryo. Ideker *et al.* (2001) have used genomic and proteomic analyses to build a detailed model of galactose utilization in yeast (*Saccharomyces cerevisiae*), that shows, among other things, that galactose uptake is a nonlinear response to the state of many interacting genes and is also connected to other physiological processes, such as respiration. Other researchers have focused on identifying the large-scale structure of molecular networks rather than on the details of a particular pathway or process. Genome-wide networks of transcriptional regulation in *Escherichia coli* (Thieffry *et al.* 1998) and *S. cerevisiae* (Wagner 2002) have been known for some time, for example, and networks of protein-protein interactions have been assessed as well (Uetz *et al.* 2000). However, much of this work detailing the properties of gene and molecular networks has yet to be examined for its influence on predictive skill within the framework of quantitative genetics applied in plant and animal breeding and ultimately its implications for evolutionary trajectories.

Recent work (Frank 1999; Cooper *et al.* 2005; Gjuvsland *et al.* 2006; Omholt *et al.* 2000; Peccoud *et al.* 2004; Dong *et al.* 2012; Marjoram *et al.* 2014) has sought to bridge the divide between classical quantitative genetics theory, rooted primarily in explaining the statistics of a population due to the collective effects of multiple unobserved genes, and the burgeoning wealth of fine-scale data on how and where genes are regulated, the proteins produced by individual genes, the biochemical pathways in which those proteins operate, and the larger-scale network of interactions within the cell. Omholt *et al.* (2000) showed how small networks of a few genes could reproduce the statistical genetics concepts of dominance, overdominance, additivity, and epistasis. Peccoud *et al.* (2004) related quantitative genetics concepts to the parameters, variables, and performance functions of a dynamic, deterministic model of galactose metabolism in yeast. They showed through *in silico* experiments that the response to selection on the output of this network suggested classical quantitative genetics properties of both additive and non-additive genetic effects that changed in relative importance over cycles of selection. Additionally, Gjuvsland *et al.* (2006) have shown that statistical epistasis is the norm when regulatory networks control trait variation. Our aim in this paper is in the spirit of these earlier authors and is motivated by the following questions. What types of predictions can we expect to make of the phenotypic behavior of a gene network? Can an understanding of the underlying molecular and genetic architecture of traits lead to improved quantitative predictions at the organism trait level? Can we predict how to change the structure of a gene network to bring about a desired change in the phenotype of a trait?

Aspects of these questions have been investigated for some time by Kauffman and colleagues using random Boolean networks as a model system (Harris *et al.* 2002; Kauffman 1995; Kauffman *et al.* 2003; Kauffman *et al.* 2004; Kauffman 1969; Kauffman 1993). Abstracting the behavior of genes as simple ON/OFF switches ignores some of the fine detail about how genes operate in real organisms. Nevertheless, the Boolean approximation has been extremely useful for understanding the properties of large-scale network systems (Bornholdt 2005) and for quantitatively predicting the behavior of networks with a small number of genes (Albert and Othmer 2003; Li *et al.* 2004). Indeed, representing a natural system by a simpler but useful predictive model is widely used in modeling biological systems (Hammer *et al.* 2006; Hammer *et al.* 2019).

A striking feature of random Boolean networks is that, despite their simplicity, they are able to produce a range of dynamical behavior, usually categorized as ordered, complex, and chaotic (Kauffman 1993). As an example, Figure 1 shows the dynamical behavior of a small Boolean network as a function of time starting from random initial conditions. In the top panel the network quickly settles into a fixed point attractor (Ott 2002). That is, the ON/OFF state of all nodes is fixed in time after about 10 time-steps. The middle panel shows a network that falls into a periodic cycle of ON/OFF states after about 330 time-steps while the bottom panel shows no regularity in the ON/OFF states of the system over the 500 time-steps we examined. Fundamentally, the goals of this paper are to progress our understanding of why some networks are ordered and others are chaotic and to determine if this behavior can be predicted for specific networks suggested from biology.

**Figure 1.**
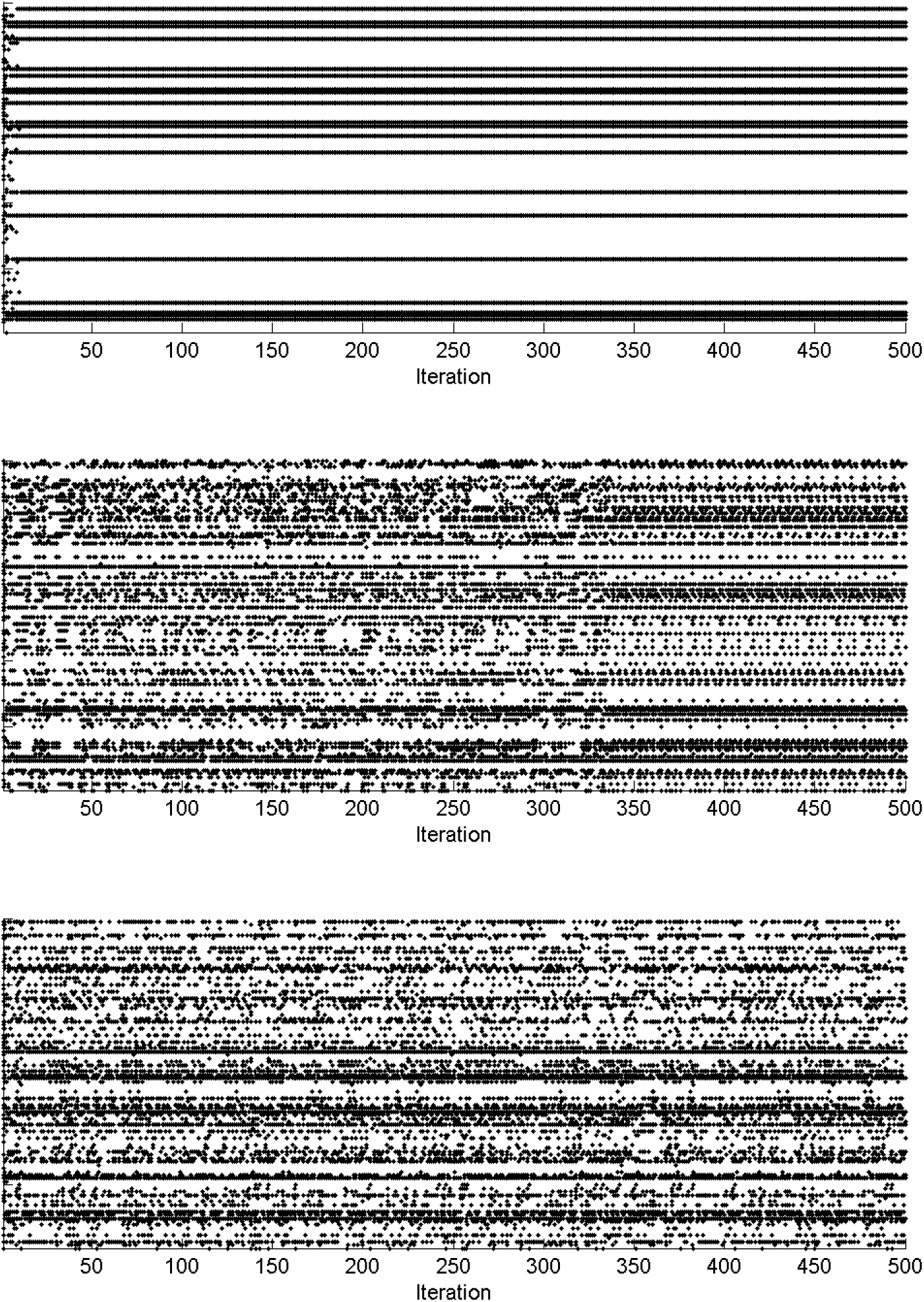
Output from an *N* = 100, *K* = 3, *P* = 0.75 random Boolean network that shows (top) ordered, (middle) complex, and (bottom) chaotic dynamics. Network nodes are listed sequentially along the y-axis and time runs along the x-axis. A white (black) cell for a node indicates that the node is OFF (ON) at that time point. *N* is the number of nodes in the network, *K* is the number of input connections for each node, and *P* is the bias probability toward being in the ON state, also referred to as a canalyzing function (Harris *et al.* 2002; Kauffman 1993). See the Materials and Methods section for further explanation.

Although characterization of the network dynamics as ordered and chaotic is qualitative, a formal mathematical definition of this transition between order and chaos has been developed (Derrida and Pomeau 1986). As such, the ordered or chaotic nature of a network may be regarded as a trait phenotype that can be measured in practice and compared to predictions based on theory. Recent papers have empirically (Harris *et al.* 2002; Shmulevich *et al.* 2005) and theoretically (Aldana and Cluzel 2003; Kauffman *et al.* 2003; Kauffman *et al.* 2004; Shmulevich *et al.* 2003) explored the importance and influence of network topology and regulatory rules on this network phenotype and its robustness to perturbation.

Ultimately, we seek to place the specific structures of real biological networks within the range of theoretical networks that can be examined. In this paper we focus our investigations on network topologies drawn from the metabolic reactions of real microorganisms and attempt to predict the phenotypes of these networks using existing theory. We approximate the system dynamics as Boolean given its usefulness and computational tractability for networks of the size we explore. The phenotype we employ is an oft-used measurement of the amount of complexity in Boolean networks: the length of attractor periods. This investigation is a step toward the larger goal of building a theory that relates the structure and function of gene networks to quantitative genetics so that the practicality of manipulating gene networks to realize improved trait phenotypes can be understood.

## MATERIALS AND METHODS

### Boolean networks

Boolean network models consider a gene network as a set of *N* genes (also referred to as network nodes) whose state may be either ON or OFF at any time *t* (Bagley and Glass 1996; Kauffman 1969; Kauffman 1993). In our formulation the state of each gene *n_i_* at time *t* + 1 was regulated by the state at time *t* of *k_i_* other genes via the transition equation 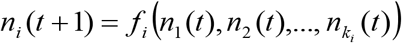. The Boolean update rule *f_i_* was selected at random, subject to the condition that its output states at time *t* + 1 are biased toward the ON (OFF) state with probability *P* (1-*P*); a network with equal probabilities of being ON or OFF at time *t* + 1 has *P* = 0.5.

Numerical simulations of random Boolean networks were generated with *N* = 100, 200, 500, or 1000. The *k_i_* input connections were assigned uniformly at random among the *N* genes such that the average connectivity 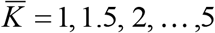. The bias probability *P* was varied from 0.5 to 0.975 in increments of 0.025. For each configuration of *N*, *K*, and *P* we created 100 random instances of a network. In the *N* = 100, 200, and 500 networks we ran 1000 separate random initial conditions. For the *N* = 1000 networks computational limitations necessitated using only 50 initial conditions. For each network, the transition equation was iterated until an attractor was found or until 100,000 time-steps were reached.

In addition to the set of random Boolean networks described above, we conducted numerical simulations of networks where the network topology was drawn from real microorganisms (see below). The network size *N* and the *k_i_* input connections for each network node within the microorganism networks were specified from metabolic reaction data for each microorganism. As with the random Boolean networks, within the microorganism networks the *P* applied for the Boolean update rule *f_i_* was varied from 0.5 to 0.975 in increments of 0.025 and up to 100,000 time-steps were simulated. Again, as with the random Boolean networks computational constraints limited the number of initial conditions, we could consider to 50 for the microorganism networks.

### Phenotypes

The state space of Boolean networks is large but finite and organized into basins of attraction (Wuensche 1998), which are the set of states that repeat sequentially when the transition equation is applied. Throughout, we refer to the repeating cycle of states as an attractor. The length of time it takes for an attractor to repeat one cycle is its period, or cycle length; we collect the period on our Boolean networks as a measure of the complexity of the network. In Figure 1, for example, the simulated network ultimately reaches an attractor with a period of 1 in the top panel and a period of 16 in the middle panel. The period length of the attractor in the last panel is undefined from the available information because over the 500 time-steps plotted no attractor was found. We adopt the convention that the attractor length in these cases is set to the maximum number of time-steps considered (*i.e*., 100,000).

A transition between ordered and chaotic system dynamics has been theoretically derived as

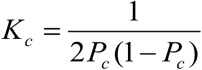

where *K_c_* is the number of input connections to each gene along the critical line that defines the transition and *P_c_* is the corresponding bias probability (Derrida and Pomeau 1986). Numerical simulation studies have previously used the length of attractor periods as a metric for characterizing the dynamics of Boolean networks (Bagley and Glass 1996; Bastolla and Parisi 1996; Bastolla and Parisi 1997; Fox and Hill 2001; Kauffman 1993). In addition to measuring the length of attractor periods a number of other network statistics were also measured.

### Network statistics

We measured several standard statistics on the microorganism networks used in this paper. The average clustering coefficient and mean shortest path between nodes were calculated using common definitions (Newman 2003b). The degree correlation between the input and output connections for all network nodes was calculated using the method outlined by Newman (2003a). A comprehensive review of each of these metrics and their properties for various graph architectures is provided by Newman (2003b).

Additionally, the distribution of the number of connections per node, referred to as the degree distribution, has often been reported for biological networks (Jeong *et al.* 2000; Lee *et al.* 2008; Lee *et al.* 2002; Newman 2003b; Thieffry *et al.* 1998; Wagner and Fell 2001). Following Stumpf *et al.* (2005) we determined the maximum likelihood fit of several standard probability distributions to the microorganism degree distributions. The Aikake information criterion (AIC)

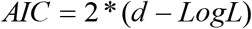

was then used as a statistically formal means to distinguish which of the model distributions best represents the degree distribution data for the microorganism network. The variable *d* is the number of free parameters required to define the model distribution and *LogL* is the log-likelihood of the maximum likelihood model for the probability distribution function. In this paper we fit the Poisson, exponential, lognormal, stretched exponential and power law distributions to the degree distributions of the microorganism networks. Details of the form of each of these distributions and their log-likelihood functions appear in the Appendix.

### Microorganism data

Metabolic reaction data were taken from the MetaCyc database of metabolic pathways and enzymes (Caspi *et al.* 2006; Caspi *et al.* 2007; Karp *et al.* 2005) for the 15 microorganisms listed in Table 1. The *Nostoc sp.* (Nostoc), *Rhodopseudomonas palustris* (Rhodopseudomonas), *Chlorobium tepidum* (Chlorobium), *Synechocystis sp.* (Synechocystis), *Deinococcus radiodurans* (Deinococcus), *Pyrobaculum aerophilum* (Pyrobaculum), *Lactobacillus plantarum* (Lactobacillus), *Methanococcus jannaschii* (Methanococcus), *Archaeoglobus fulgidus* (Archaeoglobus), *Thermotoga maritima* (Thermotoga), *Bradyrhizobium japonicum* (Bradyrhizobium), *Mesorhizobium loti* (Mesorhizobium), *Aquifex aeolicus* (Aquifex), *Desulfovibrio vulgaris* Hildenborough (Desulfovibrio), and *Nitrosomonas europaea* (Nitrosomonas) Tier III databases were produced as a collaboration between the group of Peter D. Karp at SRI International and the group of Christos Ouzounis at the European Bioinformatics Institute. The specific authors of these organism databases are Pallavi Kaipa, Peter D. Karp, and Alexander Shearer of SRI International.

**Table 1.**
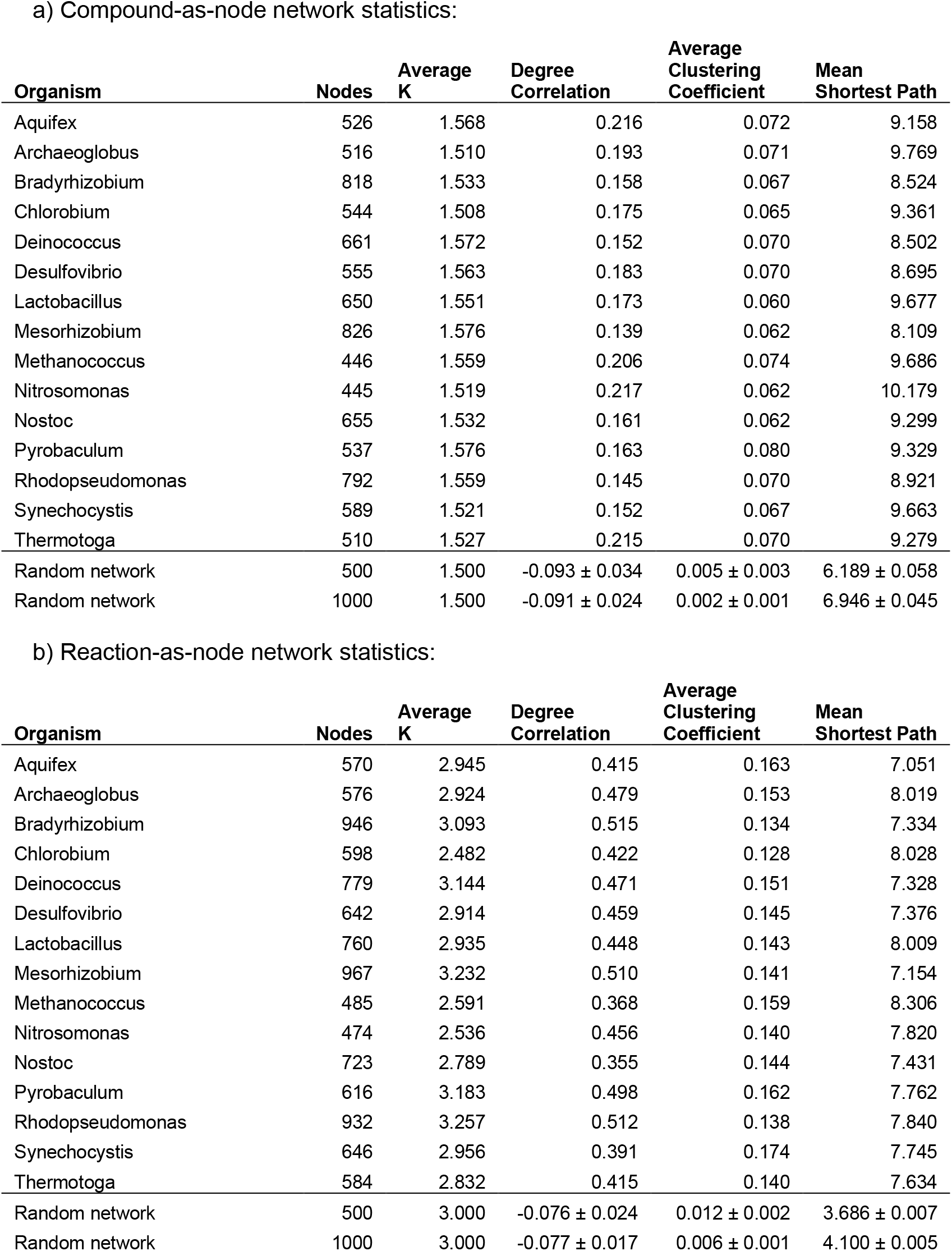
The statistical properties of microorganism network topologies when the network is calculated a) with compounds as nodes and b) with reactions as nodes. Two representative randomly constructed networks are provided for reference. For the random networks the statistical properties are estimated from 1000 randomly generated graphs.

| Organism | Nodes | Average K | Degree Correlation | Average Clustering Coefficient | Mean Shortest Path |
| --- | --- | --- | --- | --- | --- |
| Aquifex | 526 | 1.568 | 0.216 | 0.072 | 9.158 |
| Archaeoglobus | 516 | 1.510 | 0.193 | 0.071 | 9.769 |
| Bradyrhizobium | 818 | 1.533 | 0.158 | 0.067 | 8.524 |
| Chlorobium | 544 | 1.508 | 0.175 | 0.065 | 9.361 |
| Deinococcus | 661 | 1.572 | 0.152 | 0.070 | 8.502 |
| Desulfovibrio | 555 | 1.563 | 0.183 | 0.070 | 8.695 |
| Lactobacillus | 650 | 1.551 | 0.173 | 0.060 | 9.677 |
| Mesorhizobium | 826 | 1.576 | 0.139 | 0.062 | 8.109 |
| Methanococcus | 446 | 1.559 | 0.206 | 0.074 | 9.686 |
| Nitrosomonas | 445 | 1.519 | 0.217 | 0.062 | 10.179 |
| Nostoc | 655 | 1.532 | 0.161 | 0.062 | 9.299 |
| Pyrobaculum | 537 | 1.576 | 0.163 | 0.080 | 9.329 |
| Rhodopseudomonas | 792 | 1.559 | 0.145 | 0.070 | 8.921 |
| Synechocystis | 589 | 1.521 | 0.152 | 0.067 | 9.663 |
| Thermotoga | 510 | 1.527 | 0.215 | 0.070 | 9.279 |
| Random network | 500 | 1.500 | -0.093 ± 0.034 | 0.005 ± 0.003 | 6.189 ± 0.058 |
| Random network | 1000 | 1.500 | -0.091 ± 0.024 | 0.002 ± 0.001 | 6.946 ± 0.045 |

b) Reaction-as-node network statistics:
| Organism | Nodes | Average K | Degree Correlation | Average Clustering Coefficient | Mean Shortest Path |
| --- | --- | --- | --- | --- | --- |
| Aquifex | 570 | 2.945 | 0.415 | 0.163 | 7.051 |
| Archaeoglobus | 576 | 2.924 | 0.479 | 0.153 | 8.019 |
| Bradyrhizobium | 946 | 3.093 | 0.515 | 0.134 | 7.334 |
| Chlorobium | 598 | 2.482 | 0.422 | 0.128 | 8.028 |
| Deinococcus | 779 | 3.144 | 0.471 | 0.151 | 7.328 |
| Desulfovibrio | 642 | 2.914 | 0.459 | 0.145 | 7.376 |
| Lactobacillus | 760 | 2.935 | 0.448 | 0.143 | 8.009 |
| Mesorhizobium | 967 | 3.232 | 0.510 | 0.141 | 7.154 |
| Methanococcus | 485 | 2.591 | 0.368 | 0.159 | 8.306 |
| Nitrosomonas | 474 | 2.536 | 0.456 | 0.140 | 7.820 |
| Nostoc | 723 | 2.789 | 0.355 | 0.144 | 7.431 |
| Pyrobaculum | 616 | 3.183 | 0.498 | 0.162 | 7.762 |
| Rhodopseudomonas | 932 | 3.257 | 0.512 | 0.138 | 7.840 |
| Synechocystis | 646 | 2.956 | 0.391 | 0.174 | 7.745 |
| Thermotoga | 584 | 2.832 | 0.415 | 0.140 | 7.634 |
| Random network | 500 | 3.000 | -0.076 ± 0.024 | 0.012 ± 0.002 | 3.686 ± 0.007 |
| Random network | 1000 | 3.000 | -0.077 ± 0.017 | 0.006 ± 0.001 | 4.100 ± 0.005 |

Networks of the metabolic reaction data were constructed in two separate ways (Wagner and Fell 2001). First, chemical compounds were defined as nodes of the network and edges as metabolic reactions. An edge was drawn between two nodes if the chemicals were present in the same reaction. In the second formulation the metabolic reactions are defined as network nodes and the chemical compounds are network edges. An edge connected two nodes if that chemical compound was present in both reactions. Networks constructed in these two ways for two different organisms are shown in Figure 2. The layout of the graph was created using custom software that implements the Kamada-Kawai algorithm (Kamada and Kawai 1989) for optimally placing graph nodes and edges. In general, for each microorganism the reaction-as-node networks had a larger *N* and 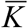 when compared to the compound-as-node networks (Table 1).

**Figure 2.**
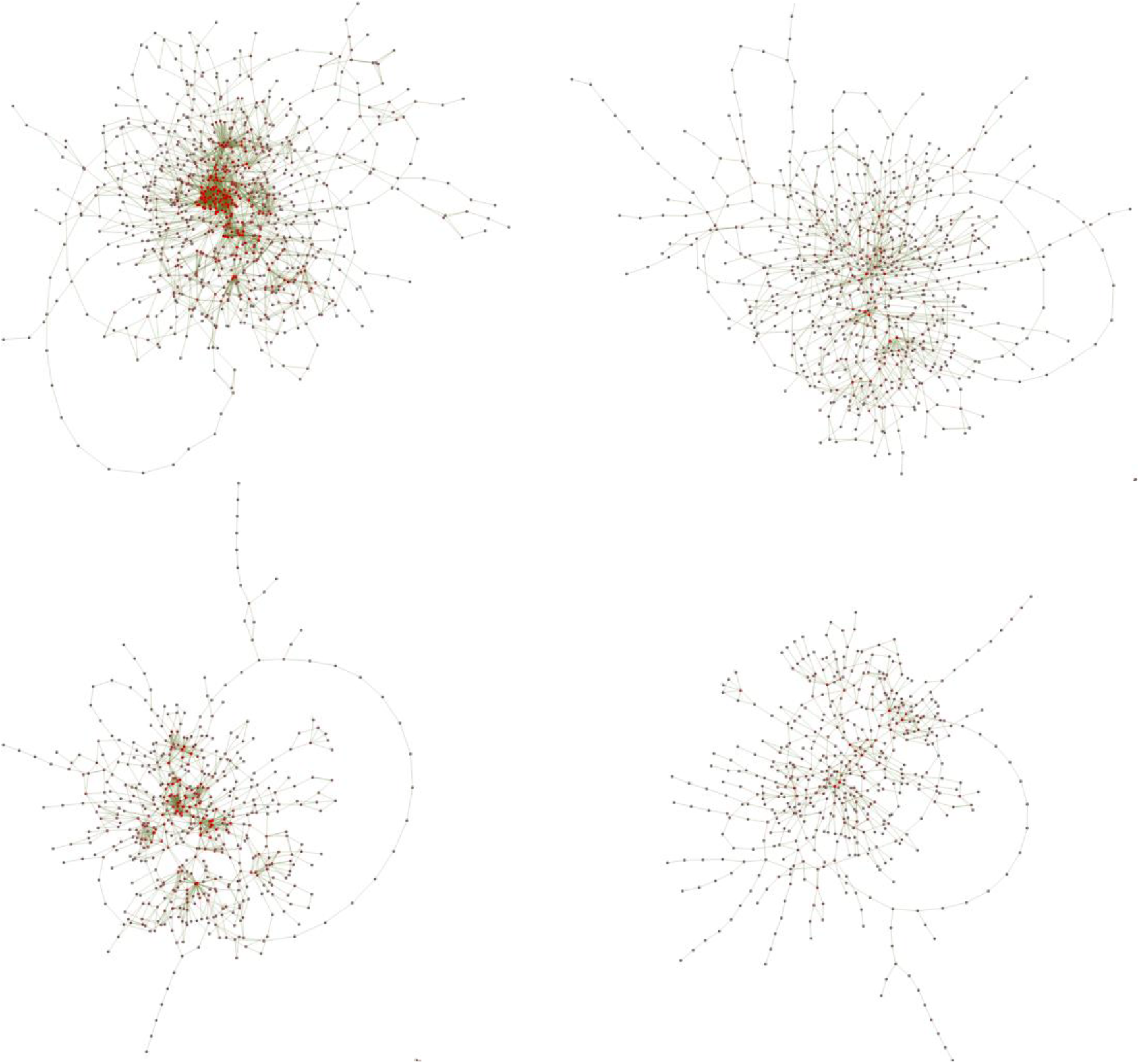
Diagrams of metabolic networks drawn from the microorganisms mesorhizobium (top) and chlorobium (bottom) when the definition of a network node is made on the chemical reactions (left) or chemical compounds (right). See text for details.

## RESULTS

Theory indicates that a transition from ordered dynamics to chaotic dynamical behavior will occur for Boolean gene networks in the limit of large *N* and given a specific *K* (Derrida and Pomeau 1986). Additionally, theory indicates that Boolean gene networks with *K* < 2 will always exhibit ordered dynamics, i.e. short attractor periods. Figure 3 shows the theoretical transition line between ordered and chaotic dynamical behavior along with a heat map representing the ensemble average attractor periods for the *K-P* state space. The coloring scheme maps short average attractor periods to blue and long average attractor periods to red. For the levels of *N* explored in this paper we find that a transition from ordered to chaotic dynamics does occur in numerical simulations, that the shape of this transition zone mimics that of the curve predicted from theory, and that networks with *K* < 2 always have short attractor periods (Fig. 3). Interestingly, however, the transition from order to chaos depends on the size *N* of the underlying network. For *N* = 100 and *P* = 0.5, for example, the average length of attractor periods is small up to *K* = 3.5, after which it increases dramatically, reaching a maximum value of 100,000 at *K* = 5. As *N* is increased the portion of Fig. 3 exhibiting long average attractor periods, and hence chaotic dynamics, gradually increases. By *N* = 1000 the theoretical order-chaos transition curve reasonably approximates the numerical simulation results.

**Figure 3.**
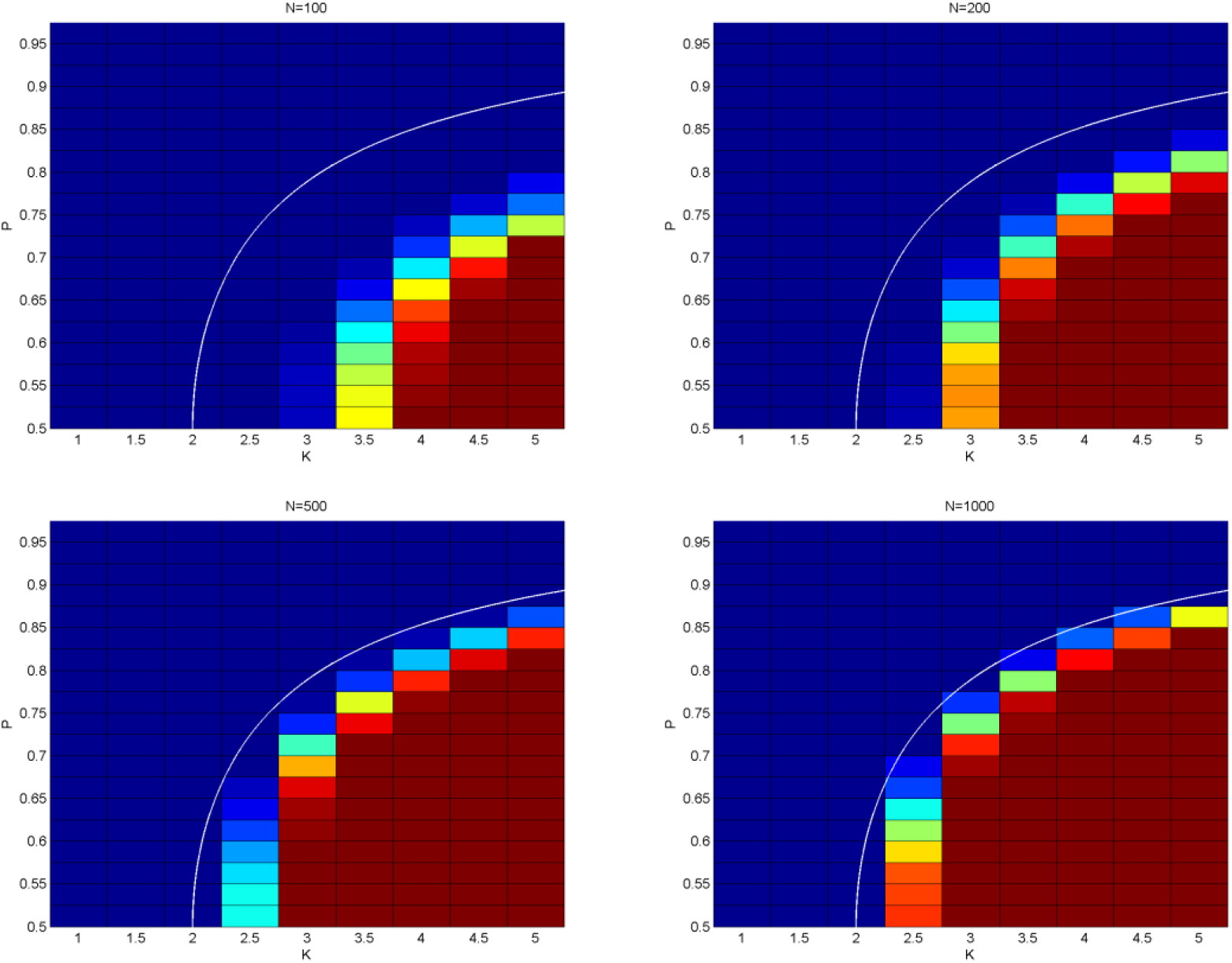
The average attractor length measured on random Boolean networks for *N* = 100, 200, 500, and 1000 at a range of *K* and *P*. The cells in each graph are color-coded based on the attractor length from a minimum of 1 (blue) to 100,000 (red). The transition from short to long attractor periods predicted by theory for large *N*, also known as the transition from ordered to chaotic dynamics, is shown as the thin white line by applying the equation given by Derrida and Pomeau (1986).

As with the random Boolean networks, a transition from ordered to chaotic dynamics was demonstrated for the networks drawn from real microorganism metabolic data (Table 1). Table 2 shows the average period as a function of *P* for the four microorganism networks shown in Figure 2. Networks with higher *K*, such as when the reactions are defined to be the network nodes, have longer periods than networks with lower *K*. In addition, the networks with higher *N* show longer average periods, consistent with the expected properties of the random Boolean networks shown in Fig. 3. The attractor phenotypes measured on the microorganism networks, however, do show differences with those measured on random Boolean networks. As an example, Fig. 4 plots the average length of attractor periods for three microorganism networks with *N* ~ 500 and *K* ~ 2.5 and that of a random Boolean network with *N* = 500 and *K* = 2.5. The length of attractor periods for the microorganism networks are short for *P* > 0.75, high for *P* < 0.6, and highly variable in the range 0.6 < *P* < 0.75. The flattening of the average attractor period at a maximum value of 100,000 arises because we imposed an upper limit of 100,000 time-steps in our simulations; if a higher number of time-steps could have been simulated we expect to see variation in the average attractor length for *P* < 0.6. The random Boolean networks, in contrast, transition from long to short average periods in the range 0.55 < *P* < 0.675. Additionally, the average period of the random graphs is much smaller than that of the microorganism networks when *P* < 0.725. At higher levels of *P* the dynamics are so strongly ordered that the average periods of the microorganism and random networks are equivalent. The differences between the average periods of random Boolean networks and the microorganism networks are not dependent on the specific microorganisms shown in Fig. 4; choosing a different level of *N* and *K* yields a similar comparison between random Boolean networks and the microorganism networks (data not shown).

**Table 2.** The average attractor period estimated from numerical simulations of the four microorganism networks shown in Figure 2 for varying levels of *P*. The size *N*, average *K*, and statistical properties for the microorganism networks are shown in Table 1.

| **P** | **Mesorhizobium** | **Chlorobium** | **Mesorhizobium** | **Chlorobium** |
| --- | --- | --- | --- | --- |
|  | **Reactions as Nodes** | **Reactions as Nodes** | **Compounds as Nodes** | **Compounds as Nodes** |
| 0.500 | 100,000 | 82,571 | 16,843 | 167 |
| 0.525 | 100,000 | 82,533 | 13,603 | 171 |
| 0.550 | 100,000 | 83,306 | 10,138 | 217 |
| 0.575 | 100,000 | 84,434 | 8,168 | 136 |
| 0.600 | 100,000 | 74,111 | 6,745 | 136 |
| 0.625 | 100,000 | 58,682 | 3,179 | 97 |
| 0.650 | 100,000 | 46,196 | 3,291 | 92 |
| 0.675 | 100,000 | 29,072 | 759 | 65 |
| 0.700 | 100,000 | 10,849 | 344 | 35 |
| 0.725 | 100,000 | 5,505 | 138 | 32 |
| 0.750 | 99,880 | 1,228 | 83 | 26 |
| 0.775 | 82,634 | 189 | 32 | 20 |
| 0.800 | 39,819 | 97 | 35 | 13 |
| 0.825 | 9,107 | 43 | 15 | 11 |
| 0.850 | 583 | 19 | 15 | 7 |
| 0.875 | 75 | 12 | 8 | 5 |
| 0.900 | 18 | 11 | 4 | 4 |
| 0.925 | 11 | 6 | 3 | 2 |
| 0.950 | 5 | 3 | 2 | 2 |
| 0.975 | 2 | 1 | 1 | 1 |

**Figure 4.**
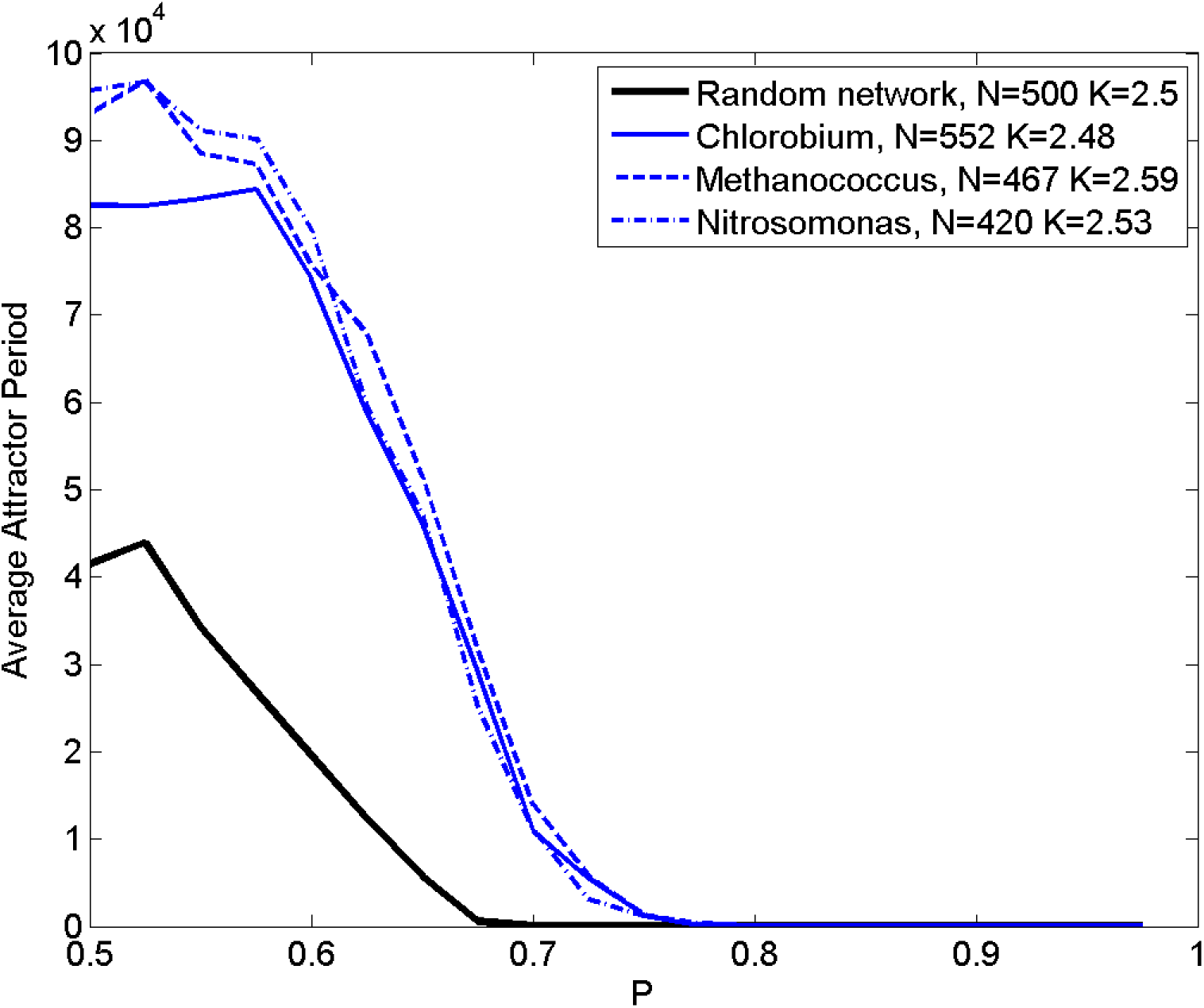
Average attractor length as a function of bias probabilities *P* for a random graph with *N* = 500 and *K* = 2.5 (thick black line) and those from the microorganism networks chlorobium (blue), methanococcus (red), and nitrosomonas (green) when defined with metabolic reactions as the network nodes and chemical compounds as the network edges.

The range of expected period phenotypes for the reaction-as-node and compound-as-node microorganism networks is shown in Table 3 for mesorhizobium and Table 4 for chlorobium, as compared to random Boolean networks with similar *N* and *K*. The range of expected period phenotypes is narrow for high *P*, regardless of *N* and *K*. The variation in expected period phenotypes is greatest in the phase transition region between ordered and chaotic dynamics for any of the networks. Also, the range of attractor period phenotypes seen in random Boolean networks with *K* = 1.5 is smaller than that seen for the microorganism networks with chemical compounds defined as the nodes of the network. A summary statistic that reflects the ability of randomly connected networks to accurately capture the dynamics of real networks is presented in Table 5. This table shows the percentage of attractor periods measured for the mesorhizobium and chlorobium microorganism networks that fall within the 99^th^ percentile of attractor periods measured from the random Boolean networks. For mesorhizobium (chlorobium), with reactions defined as network nodes, the table shows 100% agreement with the random networks for *P* < 0.75 (*P*<0.65). This agreement is a preliminary observation, however, since computational limitations prevented simulations beyond 100,000 time-steps. Networks in this part of the parameter space have had their average attractor length capped at the maximum possible value. We speculate that if the maximum time step could be increased that the percentages in columns 2 and 3 of Table 5 would be similar to those in columns 4 and 5. At higher levels of *P* the mesorhizobium networks with reactions defined as nodes often have attractor periods that fall outside the 99^th^ percentile range of the random Boolean networks. This is also a striking feature of the mesorhizobium networks with compounds defined as nodes (Table 5, column 4). The range of attractor periods measured from random Boolean networks are closer to the periods measured for the networks derived from the metabolic data of chlorobium, particularly for high values of *P* (Table 5, columns 3 and 5).

**Table 3.**
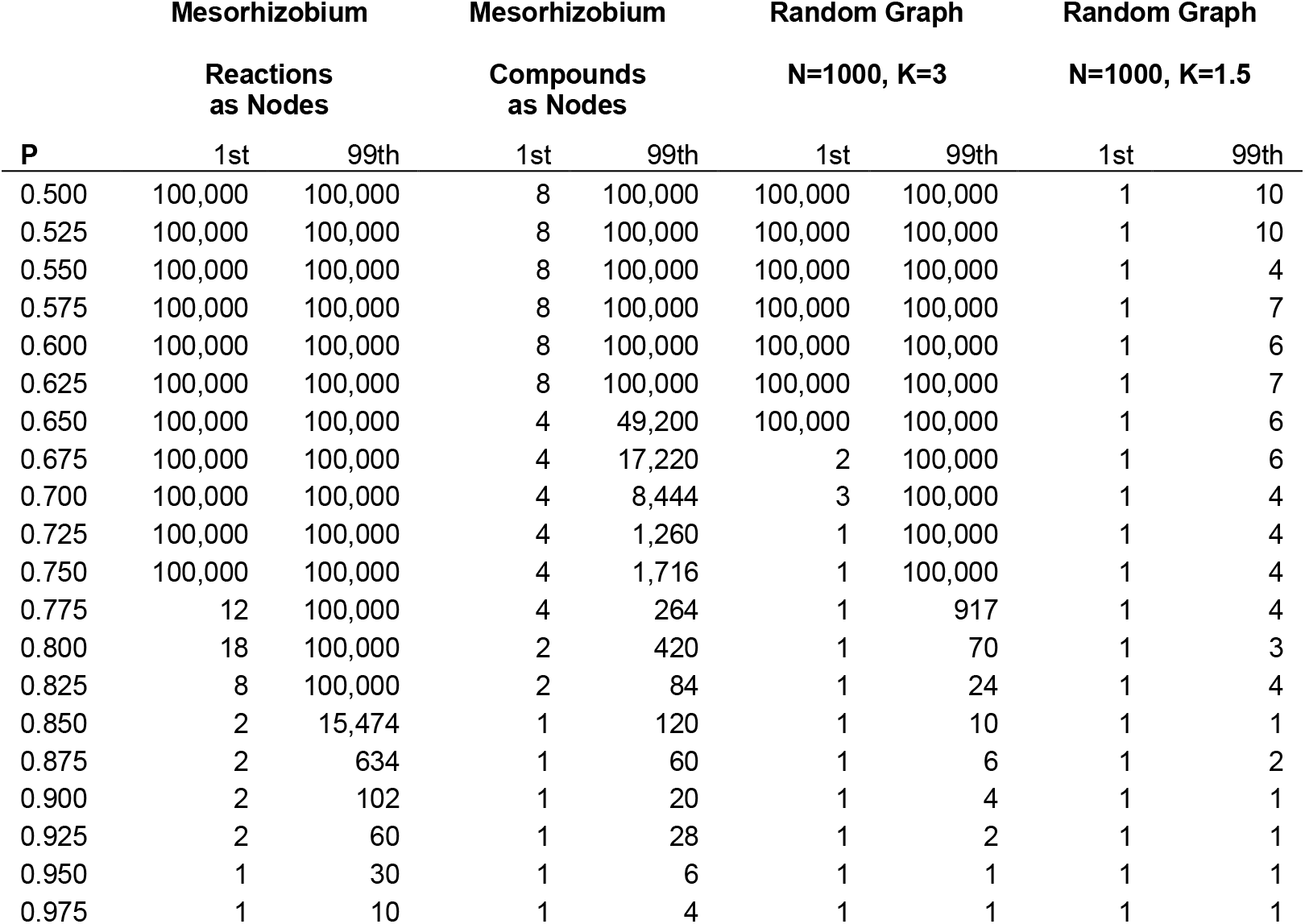
The 1^st^ and 99^th^ percentiles of attractor periods measured from numerical simulations of the mesorhizobium networks and two representative random graphs for varying levels of *P*.

**Table 4.** The 1^st^ and 99^th^ percentiles of attractor periods measured from numerical simulations of the chlorobium networks and two representative random graphs for varying levels of *P*.

| **P** | **Chlorobium** |  | **Chlorobium** |  | **Random Graph** |  | **Random Graph** |  |
| --- | --- | --- | --- | --- | --- | --- | --- | --- |
|  | **Reactions as Nodes** |  | **Compounds as Nodes** |  | **N=500, K=2.5** |  | **N=500, K=1.5** |  |
|  | 1st | 99th | 1st | 99th | 1st | 99th | 1st | 99th |
| 0.500 | 64 | 100,000 | 4 | 1,668 | 1 | 100,000 | 1 | 12 |
| 0.525 | 160 | 100,000 | 4 | 1,788 | 1 | 100,000 | 1 | 28 |
| 0.550 | 252 | 100,000 | 4 | 3,312 | 1 | 100,000 | 1 | 12 |
| 0.575 | 108 | 100,000 | 4 | 1,320 | 2 | 100,000 | 1 | 20 |
| 0.600 | 60 | 100,000 | 4 | 1,392 | 1 | 100,000 | 1 | 24 |
| 0.625 | 60 | 100,000 | 4 | 1,260 | 1 | 100,000 | 1 | 14 |
| 0.650 | 18 | 100,000 | 4 | 888 | 1 | 100,000 | 1 | 12 |
| 0.675 | 12 | 100,000 | 4 | 540 | 1 | 20,651 | 1 | 20 |
| 0.700 | 12 | 100,000 | 4 | 220 | 1 | 1,310 | 1 | 8 |
| 0.725 | 4 | 100,000 | 3 | 280 | 1 | 526 | 1 | 8 |
| 0.750 | 4 | 34,272 | 2 | 320 | 1 | 532 | 1 | 6 |
| 0.775 | 4 | 1,956 | 4 | 228 | 1 | 36 | 1 | 6 |
| 0.800 | 4 | 1,284 | 2 | 168 | 1 | 40 | 1 | 8 |
| 0.825 | 2 | 266 | 1 | 84 | 1 | 12 | 1 | 4 |
| 0.850 | 2 | 140 | 1 | 42 | 1 | 12 | 1 | 6 |
| 0.875 | 2 | 84 | 1 | 36 | 1 | 10 | 1 | 4 |
| 0.900 | 1 | 84 | 1 | 20 | 1 | 6 | 1 | 2 |
| 0.925 | 1 | 60 | 1 | 10 | 1 | 3 | 1 | 4 |
| 0.950 | 1 | 12 | 1 | 8 | 1 | 4 | 1 | 1 |
| 0.975 | 1 | 6 | 1 | 4 | 1 | 1 | 1 | 1 |

**Table 5.** The percentage of attractors for microorganism networks with periods within the 99th percentile of periods measured on random networks at similar *N* and *K*.

| **P** | **Mesorhizobium** | **Chlorobium** | **Mesorhizobium** | **Chlorobium** |
| --- | --- | --- | --- | --- |
|  | **Reactions as**<br>**Nodes** | **Reactions**<br>**as Nodes** | **Compounds as**<br>**Nodes** | **Compounds**<br>**as Nodes** |
| 0.500 | 100.0 | 100.0 | 1.3 | 16.8 |
| 0.525 | 100.0 | 100.0 | 1.4 | 42.5 |
| 0.550 | 100.0 | 100.0 | 0.1 | 21.2 |
| 0.575 | 100.0 | 100.0 | 0.8 | 27.5 |
| 0.600 | 100.0 | 100.0 | 0.6 | 32.7 |
| 0.625 | 100.0 | 100.0 | 0.4 | 29.5 |
| 0.650 | 100.0 | 100.0 | 1.6 | 32.3 |
| 0.675 | 100.0 | 65.3 | 3.6 | 42.0 |
| 0.700 | 100.0 | 57.7 | 3.6 | 26.4 |
| 0.725 | 100.0 | 60.6 | 4.4 | 32.1 |
| 0.750 | 100.0 | 78.1 | 10.8 | 32.3 |
| 0.775 | 3.3 | 41.7 | 18.5 | 37.0 |
| 0.800 | 5.9 | 59.8 | 1.1 | 63.9 |
| 0.825 | 7.5 | 40.2 | 44.8 | 58.0 |
| 0.850 | 12.4 | 67.7 | 2.0 | 73.9 |
| 0.875 | 15.0 | 59.6 | 8.3 | 81.8 |
| 0.900 | 26.4 | 64.2 | 11.0 | 37.2 |
| 0.925 | 13.0 | 44.3 | 22.2 | 95.5 |
| 0.950 | 6.4 | 83.7 | 42.6 | 51.1 |
| 0.975 | 17.8 | 44.2 | 79.5 | 81.2 |

The differences in the dynamical properties of networks drawn from microorganism data and those constructed as randomly connected graphs are driven by differences in the network topology. Table 1 lists a number of commonly calculated statistics for the microorganism networks along with values calculated for two representative random graphs. In contrast to the random graphs, the networks drawn from real microorganism data have non-zero degree correlations and average clustering coefficients. In addition, the mean shortest path on the microorganism networks is longer than expected for a random graph. Networks with compounds defined as the nodes of the network (Table 1a) have network statistics closer to those of a random graph than when chemical reactions are defined as the nodes of the network.

Since our results indicate that the topology of the microorganism networks can influence their dynamical behavior, we investigated the degree distribution of the microorganism networks. The distributions of node degree for the networks drawn from real microorganism data were fit via maximum likelihood methods to five theoretical degree distributions (Table 6). Details on the properties of the theoretical distributions are available in the Appendix. For each microorganism the theoretical distribution that best fits the underlying degree distribution is underlined. In all cases, and regardless of whether chemical reactions or compounds are defined as the nodes of the network, a lognormal distribution provided the best fit. In contrast, a random graph has a node degree distribution best represented by a Poisson function (Stumpf *et al.* 2005). Thus, the node degree distribution of the microorganism networks is different than for *NK* network models constructed as random graphs. Figure 5 shows the node degree distribution for the mesorhizobium microorganism network with reactions defined as network nodes. The observed data indicate several hundred nodes with 1 ≤ *K* ≤ 3 but also a number of nodes with *K* > 20. The Poisson distribution expected for a random graph is clearly a poor fit to the degree distribution, as evidenced by the shape of the curve in Fig. 5 and the large AIC in Table 6. Since the number of nodes with *K* = 1 is smaller than *K* = 2, the shape of the degree distribution is driven toward a lognormal fit, rather than the power law or exponential distributions that have a more linear shape for low *K* on this log-log plot. Degree distribution data for other microorganism networks show a similar pattern to that displayed in Figure 5 (data not shown). Further, we note that using the maximum likelihood estimates of the parameters of the lognormal distribution for any one microorganism network can be used to fit the parameters for the other 14 microorganisms. Doing so gives small differences in log-likelihoods, which indicates that the degree distributions of the microorganism data can be regarded as coming from the same underlying degree distribution, a result consistent with the fact that the microorganism data were collected from the same pathway genome database.

**Table 6a.**
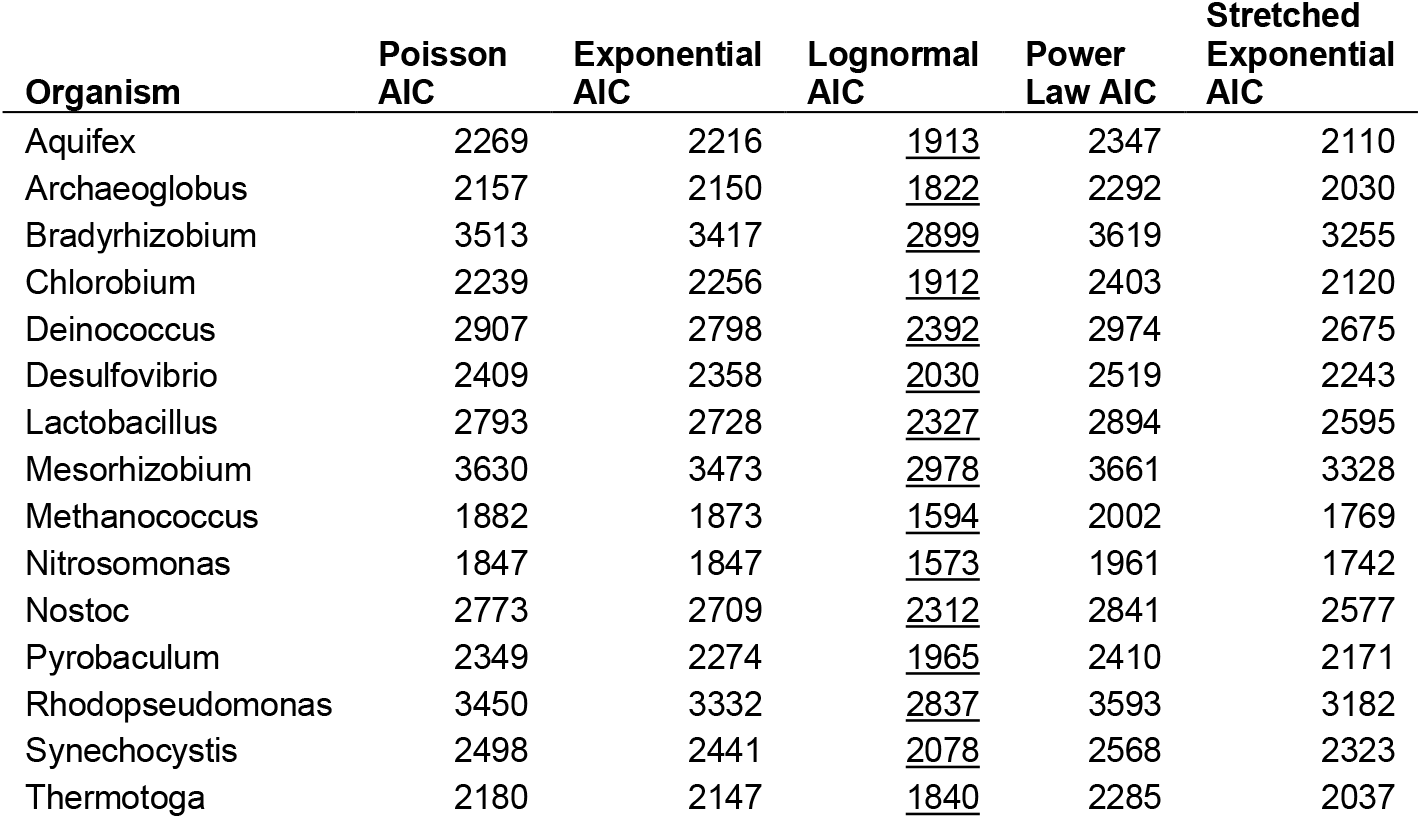
The Aikake information criterion (AIC) from statistical fits of five distributions to the node degree distribution of fifteen microorganisms. Compounds are defined to be the nodes and reactions are the edges. The model distribution which best fits the network topology of each microorganism is underlined. See the Appendix for details on the distributions and their likelihoods.

**Table 6b.**
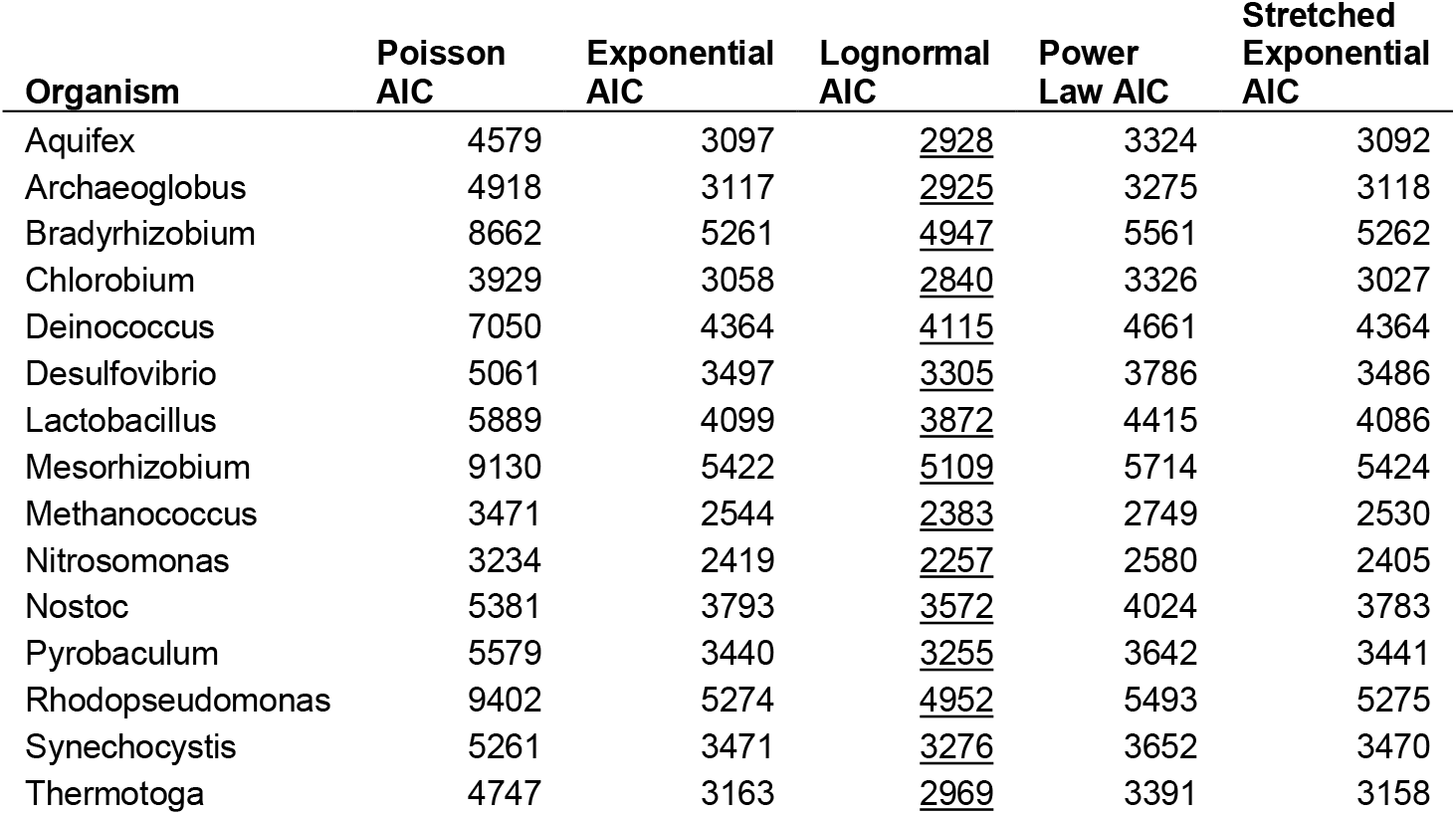
As in Table 6a, but with reactions defined to be the network nodes and compounds as the edges.

| Organism | Poisson<br>AIC | Exponential<br>AIC | Lognormal<br>AIC | Power<br>Law AIC | Stretched<br>Exponential<br>AIC |
| --- | --- | --- | --- | --- | --- |
| Aquifex | 4579 | 3097 | <u>2928</u> | 3324 | 3092 |
| Archaeoglobus | 4918 | 3117 | <u>2925</u> | 3275 | 3118 |
| Bradyrhizobium | 8662 | 5261 | <u>4947</u> | 5561 | 5262 |
| Chlorobium | 3929 | 3058 | <u>2840</u> | 3326 | 3027 |
| Deinococcus | 7050 | 4364 | <u>4115</u> | 4661 | 4364 |
| Desulfovibrio | 5061 | 3497 | <u>3305</u> | 3786 | 3486 |
| Lactobacillus | 5889 | 4099 | <u>3872</u> | 4415 | 4086 |
| Mesorhizobium | 9130 | 5422 | <u>5109</u> | 5714 | 5424 |
| Methanococcus | 3471 | 2544 | <u>2383</u> | 2749 | 2530 |
| Nitrosomonas | 3234 | 2419 | <u>2257</u> | 2580 | 2405 |
| Nostoc | 5381 | 3793 | <u>3572</u> | 4024 | 3783 |
| Pyrobaculum | 5579 | 3440 | <u>3255</u> | 3642 | 3441 |
| Rhodopseudomonas | 9402 | 5274 | <u>4952</u> | 5493 | 5275 |
| Synechocystis | 5261 | 3471 | <u>3276</u> | 3652 | 3470 |
| Thermotoga | 4747 | 3163 | <u>2969</u> | 3391 | 3158 |

**Figure 5.**
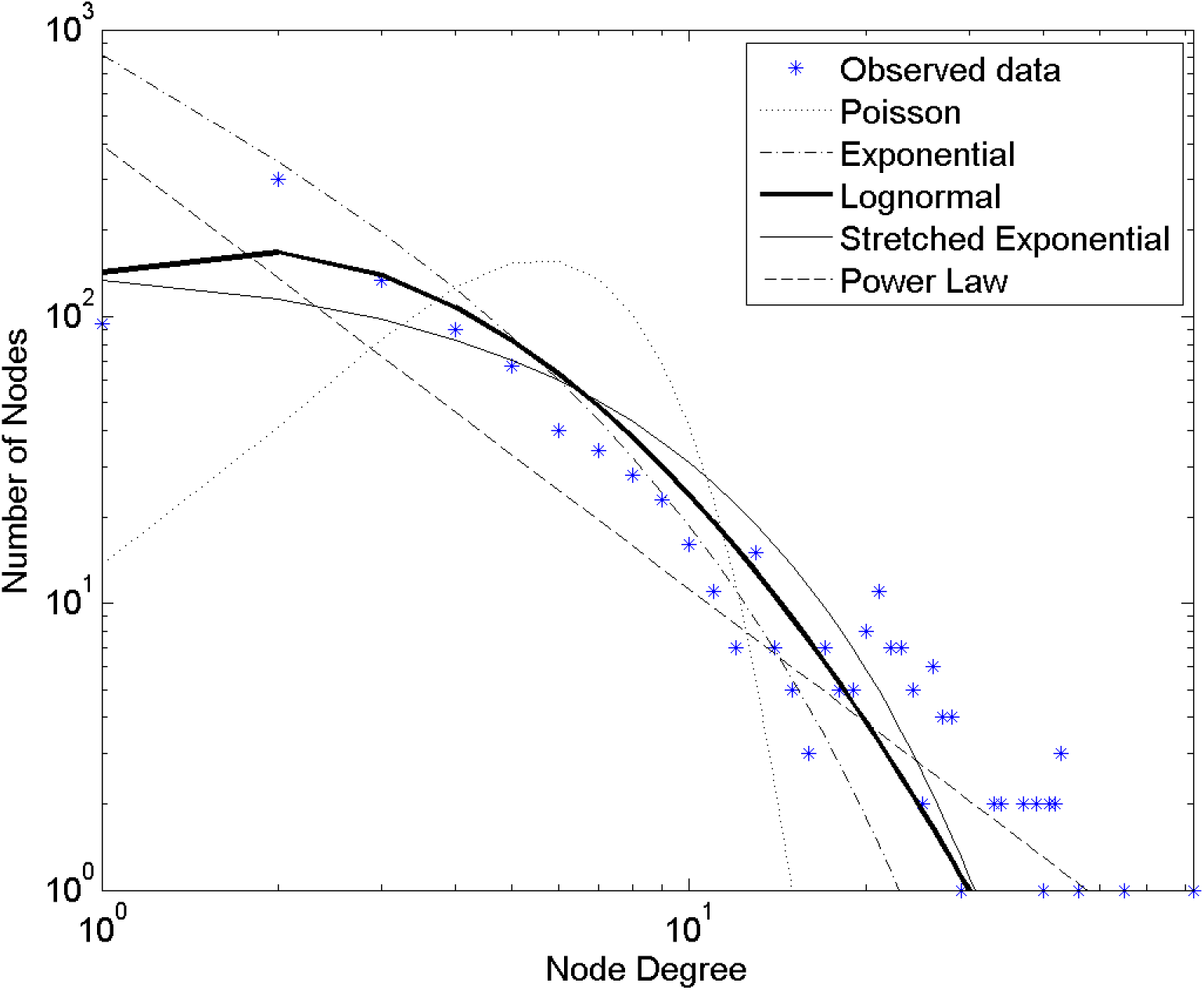
Node degree distribution for mesorhizobium with metabolic reactions defined as the network nodes and chemical compounds as the network edges. The fit of five theoretical degree distributions described in the Appendix are also plotted.

## DISCUSSION

In the many applied fields of genetics there is interest in the practicality of manipulating genes and thus gene networks to realize improved trait phenotypes (e.g., medicine, drug discovery, plant breeding, synthetic biology). Such manipulation can involve creation of *de novo* genetic variation or selection on the alleles contributing to standing genetic variation (Messina *et al.* 2011; Guo *et al.* 2014; VOSS-FELS *et al.* 2019; Wurtzel *et al.* 2019; Simmons *et al.* 2021; Powell *et al.* 2021). To achieve the expectations of this ambition requires that we can predict the behavior of networks, and their consequences for trait phenotypes, if we change some of their components, e.g., nodes or edge properties (Cooper *et al.* 2005). Previous investigations (Albert and Othmer 2003; Chaos *et al.* 2006; Peccoud *et al.* 2004) have examined the phenotypic predictability of gene networks with small *N*, whereas in this work we have focused on properties of larger networks to be representative of quantitative traits under the control of many genes (Cooper *et al.* 2005; Technow *et al.* 2021).

To develop some quantitative assessment of the feasibility of predicting the expected properties of a wide range of gene networks, we explored the predictive skill that could be expected using Boolean network theory (cf. Fig 3) as an organizing principle for an ensemble of gene network topologies. Our results reinforce the work of Derrida and Pomeau (1986) who determined that a theoretical phase transition *K*-*P* curve between “ordered” and “chaotic” network dynamics applies in the limit of large *N*. Networks of finite size do show a similar phase transition between long and short attractor periods, albeit at values of *K* and *P* different than from the theory based on asymptotically large *N*. Phenotypic predictions on random networks thus depend on knowledge of the structure of the underlying network topology, in this case through the parameter *N*. The suggestion from our results is that even random Boolean networks have phenotypes which depend crucially on the way in which the Boolean transition equation rules play out on a specific topology, especially in the phase transition region of the state space. For example, the range of observed attractor period phenotypes on a *N* = 1000, *K* = 3, *P* = 0.70 random network was between 3 and 100,000 (Table 3). To obtain such a large range in the measured attractor period phenotypes when *N* and *K* are constant across the simulations implies that knowledge of the specific rules at play in a gene network is critical to making useful predictions, a point argued at length by Wolfram (2002) and also supported by extensive empirical results obtained from studies conducted to manipulate traits of maize (Dong *et al.* 2012; Guo *et al.* 2014; Simmons *et al.* 2021). Moreover, these results call into question the utility of ensemble average properties of random Boolean networks, such as the average attractor period that we and many others have used as the trait phenotype of random Boolean networks (Bagley and Glass 1996; Bastolla and Parisi 1996; Bastolla and Parisi 1997; Fox and Hill 2001; Kauffman 1993), for predicting the phenotype of any particular real-world gene network.

To progress from the theoretical treatment to a set of network models inspired by empirical biological data on network topology, we superimposed on the theoretical space a set of network architectures drawn from a set of microorganisms available in pathway genome databases. We see this step as an attempt to identify the features of the ensemble of theoretical models relevant to our current empirical understanding of gene network architecture. The architecture of networks derived from microorganisms is clearly a non-random network topology (cf. Fig. 5, Table 1). Thus, we are encouraged to progress beyond studying network properties based solely upon ensembles of random Boolean networks. Despite this fact, the phenotypes measured on non-random networks clearly show a transition from short to long attractor periods, much like the random networks (Fig. 4). As with the random networks, however, the specific details of where in parameter space this transition happens depend strongly on the underlying network architecture. The mesorhizobium network, for example, with reactions defined as the nodes of the network shows a transition between short and long attractor periods for 0.75 ≤ *P* ≤ 0.85. In contrast, random networks of similar *N* and *K* have on average much shorter attractor periods for this range of *P* (Table 3). Additionally, even at *P* > 0.85 the distribution of phenotypes for the mesorhizobium network shows a tendency to be wider than for the random networks. In this respect, it is interesting to note that the extant microorganism network architectures that evolutionary processes have shaped are able to generate more phenotypic variability for the attractor period than the random networks. This suggests that it may prove harder to predictably manipulate the ways in which a gene responds to its input connections (i.e., altering *P*) in real networks than expected from existing random Boolean network theory. At the very least a detailed understanding of the gene network topology and regulatory rules for the networks targeted for manipulation will be required for such an undertaking.

A large body of literature has developed on calculating the so called “scale-free” nature of real networks (Farkas *et al.* 2003; Jeong *et al.* 2000; Ravasz *et al.* 2002) as well as the dynamical properties of “scale-free” networks with Boolean logic rules (Aldana and Cluzel 2003; Zhou and Lipowsky 2005). In these papers “scale-free” is taken as a pseudonym for the power law distribution (see the Appendix for details). Most of these treatments, however, do not quantitatively assess the goodness of fit of the power law distribution to the underlying data, instead relying on graphical analysis to derive a conclusion about the appropriateness of a given distribution. The graphical analysis approach is prone to bias in the estimated parameters (Goldstein *et al.* 2004) and does not provide a means to quantitatively distinguish between closely related distributions (Arita 2005). Here we statistically selected the model distribution that best fits the degree distribution of the microorganism metabolic networks (Stumpf *et al.* 2005) and found the lognormal distribution to always give the best representation of the empirical data.

Understanding how the network architecture is defined is a key to any experimental result utilizing network properties such as the degree distribution; characterizing network nodes as either metabolic reactions or compounds has a striking impact on the network topology under study and the phenotypes that are measured on that network (Tables 1 and 2). Determining the appropriate way to define the network topology depends on the question any researcher wishes to ask (Arita 2005; Wagner and Fell 2001). In this study we have tried to remain neutral on this point and instead have focused on showing how this definition can play a role in the predictions one can make.

To move beyond descriptors like “ordered” and “chaotic” to quantitative predictions of network function will require an understanding of which network structures or rules are key ingredients in determining that dynamical network behavior (e.g., Peccoud *et al.* 2004; Dong *et al.* 2012; Marjoram *et al.* 2014). We need to understand which topological characteristics of networks provide the power to understand the dynamical behavior of the system just as we need to understand how the rules embedded within these networks impact system dynamics. Ultimately such understanding of gene network properties must be examined for impact on predicting the generated variation of trait phenotypes. Our work has explored how the number of network nodes and the degree distribution of a network relate to the observed behavior of the system, i.e., the trait phenotype. We selected the attractor length of the network as a trait phenotype since this can be directly measured from the simulated ON/OFF gene expression behavior. Other trait phenotypes can easily be considered as consequences of the ON/OFF switching of genes in the network within the growth and development process of the target organism (e.g., Hammer *et al.* 2006; Peccoud *et al.* 2004; Dong *et al.* 2012; Wurtzel *et al.* 2019; Cooper *et al.* 2021; Powell *et al.* 2021). Further, we can model applications of quantitative genetic theory to predict outcomes of the selection process applied to populations of genotypes where the genetic architecture of traits is a consequence of the dynamic behavior of gene networks (Cooper *et al.* 2005; Hammer *et al.* 2006; Messina *et al.* 2011; Technow *et al.* 2021). Given our selected phenotype, we have found that: (i) the transition zone between long and short attractor periods depends on the number of network nodes for random networks (small networks behave differently than the average expectations of large *N* networks); (ii) networks derived from metabolic reaction data on microorganisms also show a transition from long to short attractor periods, but at higher values of the bias probability than in random networks with similar numbers of network nodes and average node degree, which thus suggests that empirical studies of gene networks need to account for the details of gene regulation at play in those networks to have predictive skill; (iii) the distribution of phenotypes measured on microorganism networks shows more variation than random networks when the bias probability in the Boolean rules is above 0.75; and (iv) the topological structure of networks built from microorganism metabolic reaction data is not random, being best approximated, in a statistical sense, by a lognormal distribution (suggesting limits to the insight that can be obtained about network behavior from studying the ensemble-average properties of random Boolean networks). The current study emphasizes a logical step in the direction of understanding genetic variation within a gene network context for its impact on trait phenotypic variation. The suggested step is to build from the current body of network theory based on expectations from ensembles of random networks toward an emphasis on the properties of specific empirical gene networks linked to trait phenotypes under the influence of evolutionary forces (e.g., Dong *et al.* 2012; Guo *et al.* 2014; Simmons *et al.* 2021). This shift in emphasis provides a foundation for advancing the study of quantitative genetics for gene networks that can be applied to the standing genetic variation in extant reference populations of organisms (Cooper *et al.* 2005; Marjoram *et al.* 2014; Cooper *et al.* 2021; Powell *et al.* 2021).

## APPENDIX

Properties of the five model distributions considered in this paper follow.

## Poisson

The Poisson distribution is the expected degree distribution for a large random graph. Its probability distribution is given by

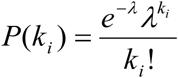

The log-likelihood of a Poisson distribution is

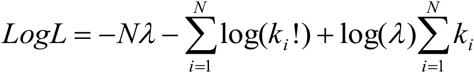

The maximum likelihood estimate of *λ* is given by

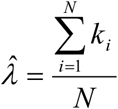

## Exponential

For the exponential distribution the probability distribution, log-likelihood function, and maximum likelihood estimate of *λ* are given by

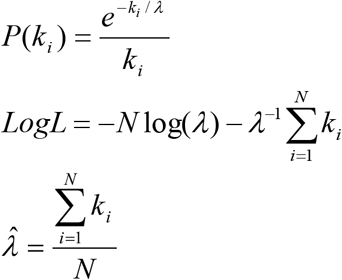

## Lognormal

For the lognormal distribution the probability distribution, log-likelihood function, and maximum likelihood estimate of *μ* and *θ* are given by

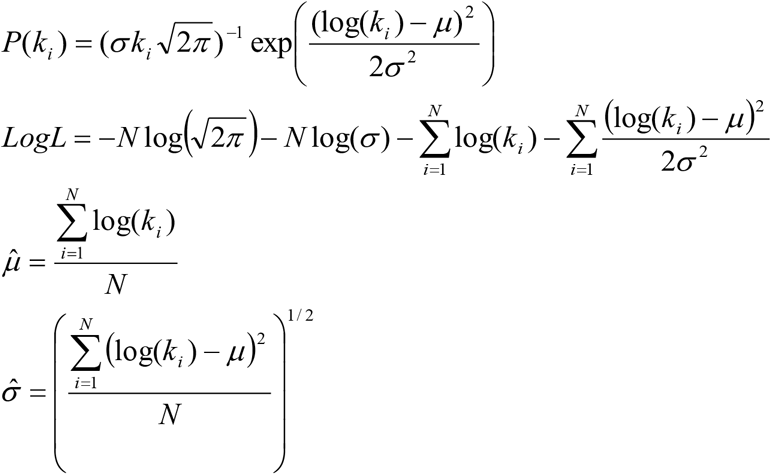

## Stretched Exponential

The probability distribution and log-likelihood function of a stretched exponential distribution are given by

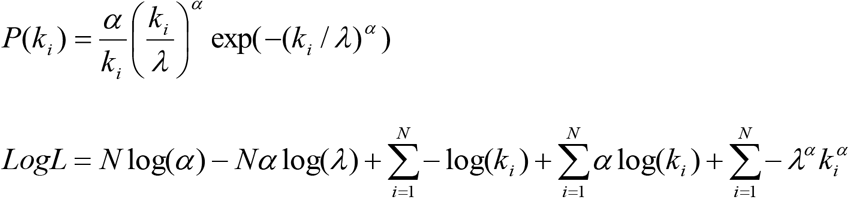

No closed form solution exists for the maximum likelihood estimates of *α* and *λ*. An approximate solution can be found by numerically determining roots of

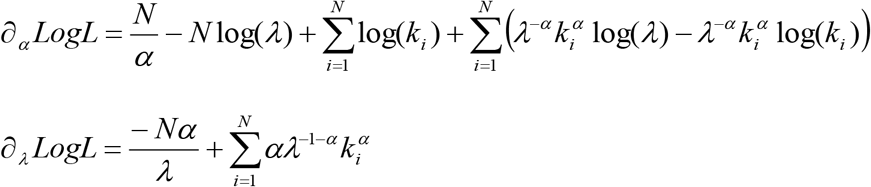

## Power Law

The power law probability distribution and log-likelihood function is given by

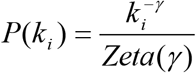

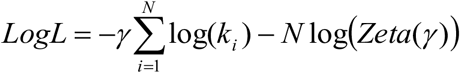

where *Zeta*(*γ*) is the Riemann zeta function. No closed form solution exists for the maximum likelihood estimate of *γ*. The approximate solution can be found by determining the point at which

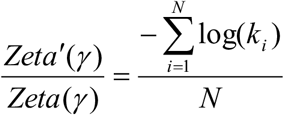

